# The influence of transcranial magnetic stimulation of the medial prefrontal cortex on emotional memory schemas

**DOI:** 10.1101/656348

**Authors:** Leonore Bovy, Ruud M.W.J. Berkers, Julia Pottkämper, Rathiga Varatheesvaran, Guillén Fernández, Indira Tendolkar, Martin Dresler

**Author notes:** Corresponding author: Leonore Bovy, Kapittelweg 29, 6525 EN, Nijmegen, The Netherlands. / /.

## Abstract

Memory bias for negative information is a critical characteristic of major depression, but the underlying neural mechanisms are largely unknown. The recently revived concept of memory schemas may shed new light on memory bias in depression: negative schemas might enhance the encoding and consolidation of negative experiences, thereby contributing to the genesis and perpetuation of depressive pathology. To investigate this relationship, we aimed to transiently perturb processing in the medial prefrontal cortex (mPFC), a core region involved in schema memory, using neuronavigated transcranial magnetic stimulation (TMS) targeting the mPFC. Forty healthy volunteers first underwent a negative mood induction to activate negative schema processing after which they received either active inhibitory (*N* = 20) or control (*N* = 20) stimulation to the mPFC. Then, all participants performed the encoding of an emotional false memory task. Recall and recognition performance was tested the following morning. Polysomnographic data was recorded continuously during the night before and after encoding. Secondary measures included sleep and mood questionnaires. We observed a significantly lower number of false recognition of negative critical lures following mPFC perturbation compared to the control group, whereas no differences in veridical memory performance were observed. These findings were supported by reaction time data. No relation between REM sleep and (false) emotional memory performance was observed. These findings support previous causal evidence for a role of the mPFC in schema memory processing and further suggest a role of the mPFC in memory bias.

## Introduction

One of the major cognitive symptoms in depression is a mood-congruent memory bias for negative information (1, 2), meaning that depressed individuals are more likely to remember negative material than those without depression (3). According to Beck’s influential cognitive model of depression, memory biases during acquisition and processing of information play an important role in the onset and maintenance of depression (4). These biases even persist after apparent recovery (5).

Despite decades of research in establishing the existence and role of negative memory biases, the neural mechanisms underlying memory bias are still elusive and have mainly been attributed to the role of amygdala (6) and its modulation of the hippocampus (7). However, the newly-revived concept of schema as applied in the cognitive neuroscience of memory (8) may shed new light on the neural mechanisms underlying memory bias in depression.

Schema-related memory processing refers to the integration of newly acquired congruent information into an existing superordinate and adaptable knowledge structure, which leads to superior encoding and consolidation efficiency (9–11). In simpler terms, new information that fits well into an existing congruent network of information (i.e. a schema), is generally remembered better (12). This benefit in learning is dependent on the schema’s activation and accessibility (13). Whereas the standard model of memory consolidation assumes an initial critical role of the medial temporal lobe (MTL) and hippocampus and a subsequent gradual integration into neocortical representations, recent literature has identified the medial prefrontal cortex (mPFC) as a key player in the immediate integration of novel memories into pre-existing knowledge structures (14, 15). Differential roles of the mPFC and the MTL in schema-dependent encoding were found, namely encoding-related activity in the mPFC increased linearly with increased congruency whereas encoding-related MTL activity decreased (16). Additionally, patients who suffer from damage to the mPFC show reduced schema representation (17, 18). These findings indicate the necessity of the mPFC for schematic memory formation. In support of the causal role of the mPFC in schema memory formation, a recent neurostimulation study (19) used transient magnetic stimulation (TMS) to perturb medial prefrontal processing before the performance of a Deese-Roedinger-McDermott (DRM) task (20). In this false memory task, participants are presented with lists of semantically related words (e.g. tired, night, pillow) that establish a schema related to a specific nonpresented semantic associate, ‘the critical lure’ (e.g. sleep). The mPFC can be inhibited by continuous theta burst (cTBS) stimulation, leading to a reduction in false recall of critical lures (19), in line with studies on mPFC lesioned patients (18). Note however that these findings are related to knowledge structures in general and it remains to be investigated how these findings transfer to emotional memory.

In depressed patients, previous studies have repeatedly shown increased amygdala activity (2, 21, 22), a structure that is often thought to play a pivotal role in the consolidation of emotional memories (23). In addition, particularly in chronic cases of depression, hippocampal function and structure shows progressive damage (24) and a reduction in size (25). Notably, increased mPFC activation is a further characteristic finding in depressed patients (26) and has been suggested as a biomarker for depression and its downregulation as a marker for therapeutic success (27, 28). In support, a double dissociation of valence-specific response in an emotional go/no-go task was found in the rostral ACC extending to the anterior medial prefrontal cortex (29). Specifically, this region was activated more strongly in response to sad words in depressed patients than to happy words, whereas this pattern was reversed in controls. An imbalance between mPFC and hippocampus, in combination with increased amygdala activity, may contribute to an increase in the recruitment of negatively toned memory schemas.

Behavioral evidence supporting this proposed mechanism comes from studies employing the DRM false memory task in depressed patients. Here, patients demonstrate a greater false memory for negative critical lures, corresponding to negative memory schema processing, in both free recall (30) and recognition (31). Similar results are found under negative mood induction conditions in healthy participants, where an induced negative mood leads to a preferential schema-like memory for negative critical lures (32–34).

Here, we propose schema memory processing as a candidate mechanism accounting for memory bias in depressed patients, who have particular strong negative schemas leading to an enhanced encoding and consolidation of negative experiences. Strong negative schemas may in fact account for the reduced tendency to process positive information even when in remission from depression (35), thereby possibly accounting for the high relapse rate in depression.

Sleep is known to play an important role in memory consolidation (36). The consolidation of both affective and schema-related memories has been associated with rapid eye moment (REM) sleep (37, 38). Furthermore, neuroimaging studies have reported increased mPFC and amygdala activity during REM sleep (39, 40). REM sleep disinhibition has been considered a biological marker of depression, which is substantiated by decreased REM latency, increased REM duration and increased REM density (41). Consequently, increased REM sleep has been suggested to increase the consolidation of negative emotional memories in depression (42). Importantly, the amount of REM sleep and prefrontal theta power during REM sleep has also been shown to predict both emotional (43) and schematic memory consolidation (37).

In sum, several lines of research point to an intimate relationship between negative memory bias, schema-related mPFC processing and sleep. However, these three components have as of yet not been integrated. To provide further evidence for this relationship, we sought to perturb schema memory processing using neuronavigated medial prefrontal stimulation, in line with a recent neurostimulation study (19). In a sample of 40 healthy participants, we used a validated experimental approach for negative mood induction (44) to activate negative schema processing as an experimental model of depressed state. We employed an adapted emotional version of the DRM false memory task to probe schema memory for both positive and negative critical lures. Lastly, we explored the relation of REM sleep to emotional and schema memory consolidation. In the preregistration of this study, we hypothesized first, that the active stimulation would diminish the number of total critical lures falsely remembered. We hypothesized, second, that the effect of the stimulation would specifically influence negative schemas due to their activation after the negative mood induction, resulting in a diminished negative memory bias in comparison to the control group. Third, we hypothesized that the amount of REM sleep and frontal theta power during REM sleep in the night between encoding and retrieval would be related to the number of critical lures that were falsely remembered.

## Methods and Materials

### Participants

Forty healthy native Dutch participants (26 females, 14 males, mean age 21.70 [± SD = 3.33], ranging between 18 and 34 years) participated in return for course credit points or monetary compensation. All participants were right-handed, had normal or corrected-to-normal vision and no history of psychiatric or neurological illness as indicated through self-reporting. The study was conducted in accordance with the Declaration of Helsinki and approved by the local medical ethical committee (CMO Region Arnhem-Nijmegen, the Netherlands with Protocol ID: NL57736.091.16) and was preregistered at Netherlands Trial Register (NTR7080/ Trial NL6893).

### Experimental procedure

The study consisted of three sessions on separate days. The first session was an intake session meant to collect baseline measurements, determine motor thresholds (MTs) and check for TMS tolerability. After providing written informed consent, participants were pre-screened on any contraindication for TMS stimulation and prepared for the TMS intervention. This consisted of head registration for neuronavigation as well as resting and active MT determination. Then, a “practice” stimulation was performed to ensure participants were comfortable with the stimulation for the following day. This practice stimulation was identical in location, duration and intensity to the actual stimulation they would receive the following day. Next, participants filled in baseline questionnaires (BDI-I, STAI-trait, NEO-PI-R, MEQ, PSQI, SMH, PANAS). Finally, the portable sleep EEG system was attached for the baseline sleep recording. Intake sessions were always scheduled in the evening (after 6 pm) and participants were instructed to not partake in any rigorous activity for the rest of the evening after EEG application.

The following day, the first experimental session took place. First, participants were prepared for the TMS session, which included head registration for neuronavigation. Then, participants filled in questionnaires (SMH and PANAS). Next, a negative mood was induced through watching several film clips. After the movie, a second PANAS was administered to probe the effectiveness of the mood induction. Immediately thereafter, the participants received either cTBS (*N* = 20) or rTMS (5 Hz; *N* = 20) stimulation for 40 seconds, targeting the mPFC. In a neighboring behavioral lab, they performed the emotional DRM encoding task. Participants were instructed to remember as many words as possible and were reminded they would be tested the following day. Finally, a third PANAS was filled in and the portable sleep EEG was again attached. The following morning, during the second experimental session, participants freely recalled as many encoded words as possible, followed by a recognition task. After completing all tasks, subjects responded to debriefing questions on the purpose of the study. See figure 1 for a schematic overview.

**Figure 1:**
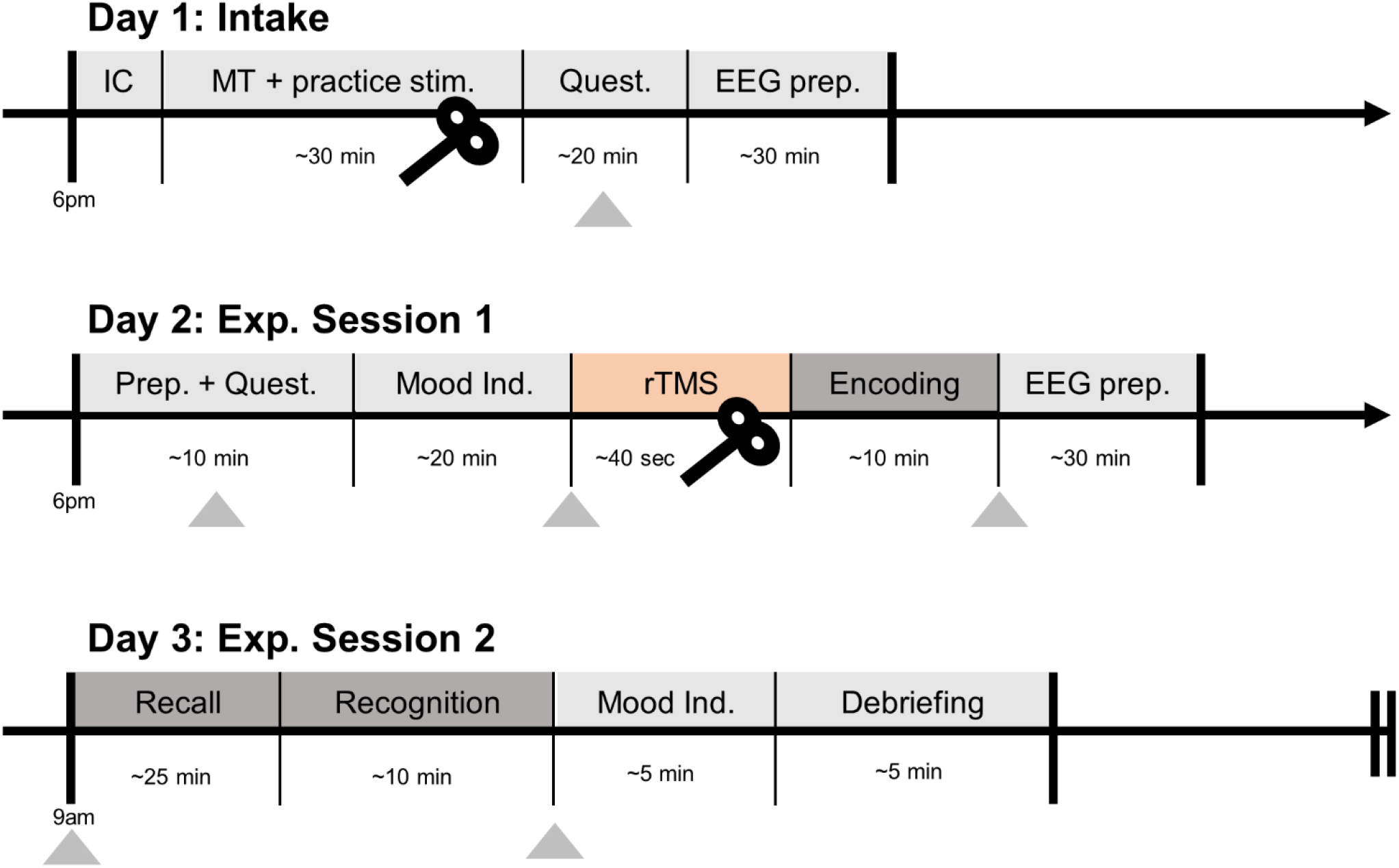
Experimental procedure. Overview of procedures during the three testing days. The grey triangles represent the administration of a PANAS questionnaire.

### False memory task

An adapted version of the well-established Deese-Roediger-McDermott (DRM) false memory task was used (20). To probe the activation of emotional schemas and measure emotional memory bias, we constructed new emotional semantically associated lists using the Dutch version of the Affective Norms for English Words (ANEW) list (45) in an initial pilot study. See supplementary materials for further details. Each list contained 10 words corresponding to one critical lure. Each word was presented twice (first in the order of strongest to weakest association with the critical lure, then immediately thereafter in semi-random order) on a computer screen for 600ms with a 125ms inter stimulus interval (ISI). Twenty lists of 10 words each were presented during encoding, corresponding to 10 positive and 10 negative lists. During the free recall, participants were asked to provide confidence ratings (1-5) for each reported word, to maximally ensure they would write down as many words as possible. The participants were not given any time limit and on average took 26.14 (± SD = 9.52) minutes. During the recognition task, three old, three new and one lure items per list were semi-randomly presented on the screen. Here, the participant had to indicate if the word was new or old, and immediately provide a confidence rating (sure, somewhat sure, unsure), within 3 seconds. To disentangle the effects of specific sleep parameters on the consolidation of false memories, encoding and retrieval sessions took place on consecutive days with a full night of EEG-recorded sleep in between. This design is in line with previous studies probing the effect of sleep on false memory processing (e.g. 46).

### TMS procedure

The active cTBS intervention and procedure was an exact copy of the study by Berkers and colleagues (2017). The cTBS stimulation protocol consisted of a total of 600 pulses administered across 40 sec, which consisted of bursts of 3 pulses at 50 Hz, and each burst was repeated at a frequency of 5Hz. The control rTMS stimulation protocol consisted of a total of 200 pulses, administered across 40 sec, at a frequency of 5Hz. Stimulation intensity was set at 80% of the active motor threshold from the tibialis anterior. See supplementary materials for further details on the TMS procedure.

### Mood induction

A sad mood was induced in all participants by presenting segments of an emotional movie (47), in order to activate negative schema memory processing, and thereby emulating a negative memory bias as seen in depressed patients (30). Two consecutive segments of the movie *Sophie’s Choice* were presented for 20 minutes (adapted from 44). See supplementary materials for participant instructions. At the end of the final experimental session, a short segment from the movie ‘Happy Feet’ was presented to induce a positive mood and to uplift any possibly lingering negative moods (44).

### Sleep recordings

Overnight polysomnographic sleep recordings were made for two consecutive nights at the participants’ home, using a 15-channel Somnoscreen™ plus (SOMNOmedics GmbH, Randersacker, Germany) portable device. Electroencephalography (EEG) measurements were recorded with gold cup electrodes at six EEG locations (F3, F4, C3, C4, O1 and O2) and were attached according to the American Academy of Sleep Medicine (AASM) rules (48). See the supplementary materials for further details on the sleep recordings.

### Analysis

All analyses were conducted in the R programming language (version 3.5.1; R Core Team, 2018). The differences between the two experimental groups were analyzed through two-tailed Welch’s t-tests for unequal variances, unless otherwise stated. For the false memory task, the hit, false alarm and critical-lure rates were calculated. A discrimination ability index (d-prime) was calculated as follows: *d*′ = *z*(*H*) − *z*(*FA*). A higher value indicates that a subject had greater ability to discriminate between studied and unstudied items. In addition, a d-prime value on the sensitivity to falsely recognizing critical lures was calculated as follows: *d*′_*criticallures*_ = *z*(*FA*_*criticallures*_) − *z*(*FA*). Here, higher values suggest a higher propensity to falsely recognize lures. In addition, we analyzed the ratio between negative and positive critical lures as a measure of emotional biased responses. Tukey’s ladder of powers was applied when data was found to be non-normal.

## Results

### Recognition results

Participants recognized old words better than chance (mean d-prime old item = 0.60, versus chance level d-prime = 0, *t*(39) = 7.57, *p* < .001). In addition, endorsement of critical lures as old studied items was also higher than chance (mean d-prime critical lures = 0.93, versus chance level d-prime = 0, *t*(39) = 7.82, *p* < .001). No ratio difference in hit rates between positive and negative items was observed for old items, *t*(39) = −0.76, *p* = 0.45 nor for critical lures, *t*(39) = 1.70, *p* = 0.10).

The d-prime scores did not differ between the two groups, *t*(38) = −0.51, *p* = .615. Furthermore, the groups did not differ in ratio of hit rates between positive and negative items for old items, rTMS: *M* = 1.00, cTBS: *M* = 0.93), after Tukey’s ladder of powers was applied due to non-normality, *t*(37.26) = 0.88, *p* = 0.38.

In accordance to our main hypotheses, the participants in the active stimulation (cTBS) group falsely remembered on average less lures (*M* = 14.10, *SD* = 3.26) than those in the control (rTMS) group (*M* = 16, *SD* = 3.26). This difference was statistically significant on a one-tailed t-test, *t*(38.00) = 1.84, *p* = 0.04, see figure 2. The d-prime scores for critical lures however did not reach statistical significance with a one-tailed t-test, *t*(34.29) = 1.57, *p* = 0.06.

**Figure 2:**
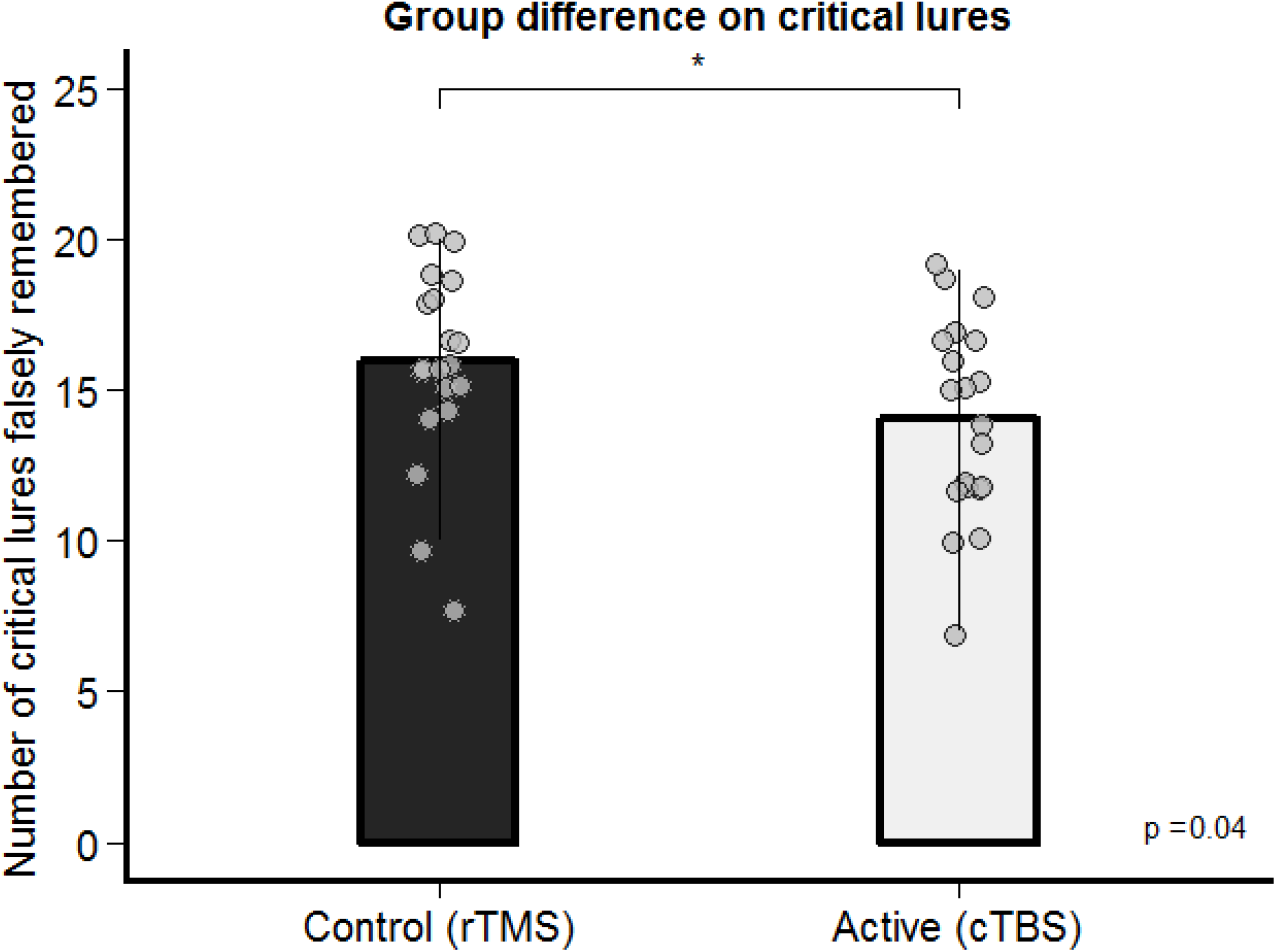
Group differences on the recognition of critical lures. A significant difference between the control and active stimulation group was found, where the active group falsely remembered on average less critical lures than the control group.

More crucially (after removal of 2 outliers that were more than 2 standard deviations from the mean in the cTBS group) the groups significantly differed in the ratio between the false alarm rates of positive and negative critical lures, *t*(34.20) = 2.27, *p* = 0.03. The cTBS group showed a more positive tendency or bias (see figure 3A) than the rTMS group (rTMS: *M* = 1.01, cTBS: *M* = 0.82). These results were supported by reaction time (RT) data, showing a similar group difference in the ratio of the valenced response times (positive/negative lures RT), *t*(33.5) = 2.87, *p* = 0.004, see figure 3B.

**Figure 3:**
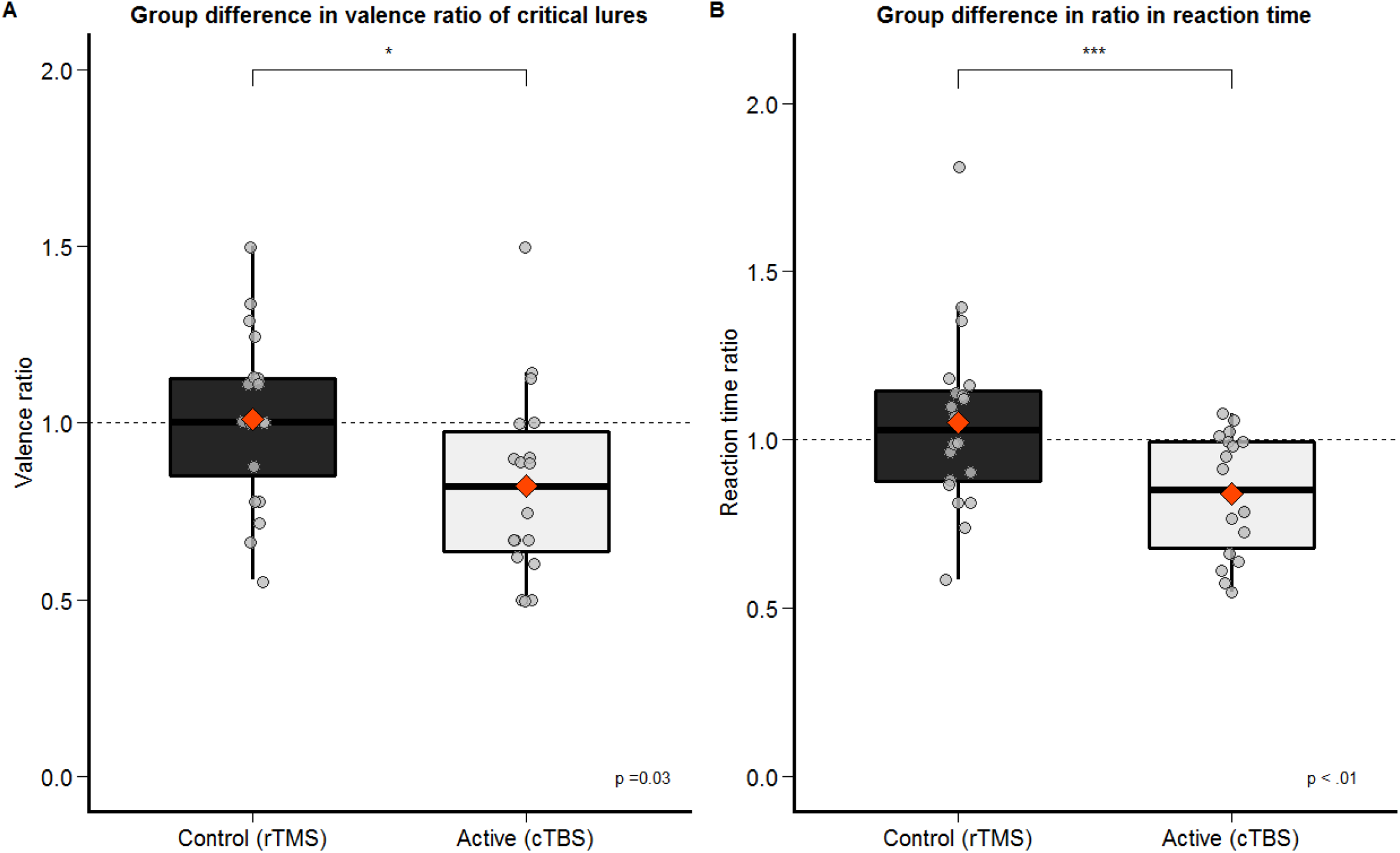
Group differences in the valence ratio and reaction time of the recognition of critical lures. A. The active group on average falsely remembered less negative critical lures relative to positive in comparison to the control group. B. The active group was on average faster in responding to positive items than negative items. The orange diamond shape represents the mean.

To explore the direction of the observed group difference in ratio scores, group differences between the false recognition of positive and negative critical lures were tested separately, revealing a decrease of false recognition of only negative critical lures in the active cTBS condition in comparison to the control condition, *t*(37.87) = 2.41, *p* = .021, see figure 4.

**Figure 4:**
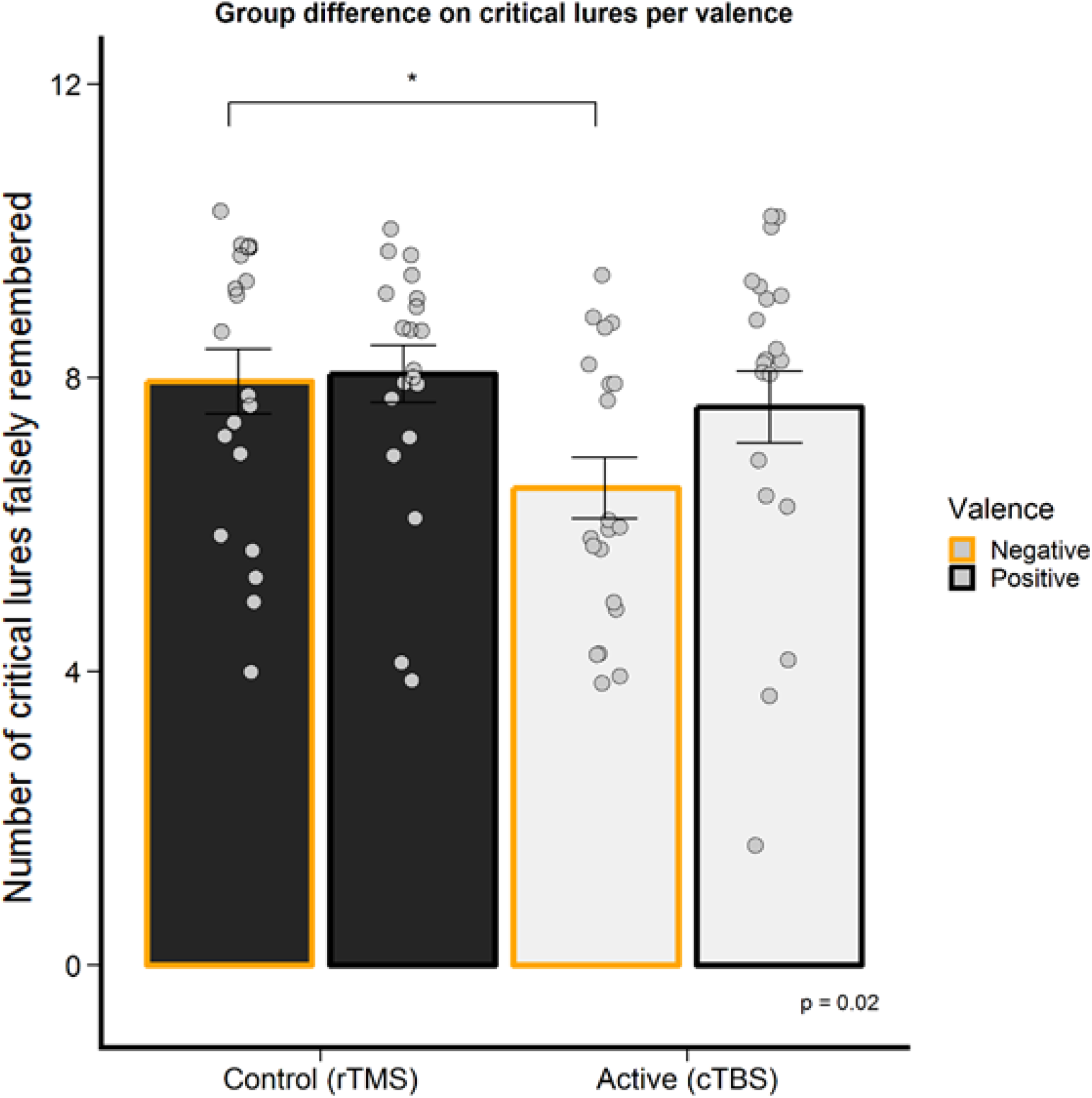
Group differences on the recognition of critical lures per valence. The main group difference was found on the negative critical lures.

### Recall results

Subjects correctly recalled on average 29.60 (SD ± 11.76) words out of all 200 studied items. No ratio difference between positive and negative items was observed for old items, *t*(39) = −1.50, *p* = 0.14. Subjects falsely recalled on average 6.22 (*SD* ± 3.19) out of 20 total lures. No ratio difference between positive and negative critical lures was found, *t*(36.97) = −1.37, *p* = 0.18. The two groups did not differ in the number of total old items remembered, *t*(38) = −0.99, *p* = .326, nor in the ratio of old items, *t*(29.49) = 0.25, *p* = 0.81. Furthermore, the groups did not differ in the total number of critical lures falsely remembered, *t*(38) = −0.84, *p* = .407, nor in the ratio of negative and positive critical lures, *t*(37) = 1.08, *p* = .285.

### Mood induction

The results of the PANAS questionnaires confirmed the effectiveness of the negative mood induction on the participants’ mood. See supplementary materials for further details.

### Sleep data

Frontal theta power during the night between encoding and retrieval was not correlated with the amount of critical lures recognized, *t*(38) = −0.10, *p* = .919, nor with the ratio between positive and negative critical lures recognized, *t*(38) = −1.38, *p* = .176. See supplementary materials for more sleep data results.

## Discussion

The present study was designed to investigate whether emotional schema memory could be reduced by transiently perturbing the mPFC using theta burst TMS. The results of the current study support the involvement of the mPFC in schema memory processing as inhibition of the mPFC with cTBS decreased false recognition in the active experimental group compared to the control group. These findings are consistent with the results of a previous TMS study (19) and with previous lesion studies (18). More importantly, after a negative mood induction, a group difference emerged in the ratio between positive and negative critical lures that were falsely recognized. Here, the active stimulation specifically decreased false recognition of negative lures. In support, the experimental groups also differed in the ratio of reaction times between negative and positive critical lure recognition, demonstrating that not only memory but also the speed of retrieval was affected by the stimulation.

The current findings show initial support to an association of mPFC-related emotional memory schemas and mood congruent memory bias. Indeed, other false memory studies using mood induction paradigms (34) have found mood-congruent effect after a negative mood induction in healthy participants. Previous meta-analytic studies have shown that healthy non-depressed individuals show a positive memory bias (49, 50) but a negative mood induction can induce a more negative memory bias (33). We propose that in our sample, the negative mood induction activated negative schemas and induced a memory bias that is mood-congruent and thus more negative. Ultimately, this resulted in a net neutral bias as we observed in the control stimulation group. In turn, we propose that the active inhibitory stimulation of the mPFC may have specifically abolished the effects of the negative mood induction by targeting the activated negative schemas, subsequently leading to a relatively more positive bias than the control group. On a neural level, the amygdala has been proposed as a mediator for mood congruent memory processing (6) and a modulator of hippocampal activity during emotional memory formation (51). Together with increased mPFC activity, increased amygdala activity and mPFC-amygdala connectivity might augment the reactivation of emotionally negative schemas. Future functional MRI studies could specifically delineate the involvement of the amygdala in connectivity to the mPFC during the encoding of emotional memory schemas.

Schemas have been considered in the cognitive theory of depression (4). Whereas in the neuroscience literature, schema is defined as a framework of knowledge that enables efficient memory storage (9), Beck’s definition of cognitive schemas includes mental representations of experiences, containing attitudes about oneself, which influence how information is being processed. These cognitive schemas in turn lead to specific memory biases in depression. We propose that these two notions are not mutually exclusive and show conceptual and neurobiological overlap. Indeed, we extend the current neurobiological model of depression (2) by suggesting that increased amygdala, hippocampus and mPFC reactivity in response to an aversive stimulus not only leads to increased ruminative thought or reinforcement of self-referential schemas, but also increased negative memory schema processing and biased memory for negative stimuli.

Previous studies reported group differences in recall (18, 19) whereas we report differences in recognition. This discrepancy could be attributed to the fact that the standard DRM task includes an immediate recall after each separate list and a recognition session immediately thereafter. Our task included a >12 hour delay between encoding and both recall and recognition. Therefore, we could dissociate between the effects of the stimulation on encoding and retrieval, but thereby increased task difficulty. A longer delay between encoding and retrieval has been shown to decrease false memory (52). This delay could have removed any ceiling effects of the recognition results as found in the previous studies which might have concealed group differences and simultaneously could have made recall too difficult to detect an effect.

There are some caveats. First, our design did not include a neutral or positive mood condition. Therefore, no specific directional effect of the mood induction can be definitively ascertained. Second, the current design included a control stimulation protocol of 5Hz applied to the same mPFC-location as the active stimulation to ensure a similar stimulation sensation in both our experimental groups. Given that one of our main hypotheses regarded responses to emotional stimuli, we aimed to exclude any group differences in mood due to potential differences in pain or discomfort. A control stimulation of 5Hz was chosen based on a previous study (53) observing no behavioral differences between a 5Hz-aPFC or cTBS-vertex control stimulation in a similar set-up as our study. However, we cannot exclude the possibility that our control stimulation did excite the cortex, thus possibly driving the group difference effect instead of the cTBS inhibition. This possibility is however not supported by the current literature on the involvement of the mPFC and memory processing. Third, the current stimulation protocol was optimized to stimulate deeper brain regions, targeting the mPFC. This however leads to additional stimulation of the areas between the stimulation coil and the target region. Berkers et al. (2017) discuss the impact of this factor and the uncertainty on the effective depth of the stimulation and will thus not be further discussed here. Lastly, to discern the directionality of the observed emotional biases compared to neutral conditions, future studies should additionally include neutral lists. In our design, no neutral lists were used due to time constraints of the encoding session, as the effect of the mood induction and brain stimulation would taper off with increased passing of time (54). Therefore, we decreased the number of lists of the encoding session by omitting the neutral condition and focused on the contrast between positive and negative stimuli to measure potential memory biases.

In summary, mPFC inhibition after negative schema activation appears to be able to specifically decrease negative schema memories. The ability to specifically target and diminish negative memories may be a valuable instrument for therapeutic purposes. Given these new findings, it would be fruitful to further explore the possibilities of manipulating emotional schemas.

## Conflict of interest

The authors declare no conflict of interest.

## Acknowledgements

Intramural funding was obtained from the Donders Center for Medical Neuroscience.

