## Supplemental materials for "The influence of transcranial magnetic stimulation of the medial prefrontal cortex on emotional memory schemas"

### Supplementary materials

#### Supplementary Methods

##### Questionnaires

In order to measure changes in mood over time during the experiment, the Positive Affect Negative Affect Schedule (PANAS; (1)) was administered repeatedly. In addition, the Becks Depression lnventory (BDI-I; (2)) was administered to measure self-reported individual differences in depressive symptoms, the State-Trait Anxiety Inventory (STAI-trait) to assess individual trait anxiety (3) and the NEO Five-Factor Inventory (NEO-FFI; (4)) to assess participants’ personality traits. Several sleep-related self-report questionnaires were administered to probe both state and trait characteristics of our participants’ sleep quality, including the Pittsburgh Sleep Quality Index (PSQI; (5)), the Morningness–eveningness questionnaire (MEQ; (6)), the Consensus Sleep Diary (CSD; core form (7)) and lastly the St. Mary’s Hospital sleep questionnaire (SMH; (8)).

###### PANAS

In this questionnaire, 20 affective items are presented to which the participant has to provide a 5-point rating in accordance with their current mood state. The items are subdivided into two categories: Positive Affect and Negative Affect. Scores per category range from 10 to 50, with higher scores representing higher levels of positive and negative affect respectively.

###### BDI

The Becks Depression lnventory (BDI-I; (2)) was administered to measure self-reported individual differences in depressive symptoms. Participants had to indicate for 21 items the extent to which these items are an accurate description of them on a scale ranging from 0 to 3 resulting in a sum score ranging between 0 and 63. The total score can be divided into four ranges, describing the severity of depressive symptoms from minimal (0-13), mild (14-19), moderate (20-28) to serious depressive symptoms (29-63). The items describe different symptoms of major depressive disorder according to the DSM-IV (APA, 1994).

###### STAI

The State-Trait Anxiety Inventory (STAI-trait) was used to assess individual trait anxiety (3). Participants had to indicate for 20 statements, how well each described their general level of anxiety on a 4-point scale. A total score ranges from 20 to 80, with higher scores representing higher trait anxiety.

###### NEO-FFI

The NEO Five-Factor Inventory (NEO-FFI; (4)) was used to assess participants’ personality traits. The NEO-FFI consists of 60 items, examining the Big Five personality traits (openness to experience, conscientiousness, extraversion, agreeableness, and neuroticism), containing 12 items per domain.

###### PSQI

To measure trait differences between participants, the Pittsburgh Sleep Quality Index (PSQI; (5)) was administered as a self-report measure of sleep quality across the previous month.

###### MEQ

The Morningness–eveningness questionnaire (MEQ; (6)) was administered to measure whether a person’s circadian rhythm (biological clock) produces peak alertness in the morning, in the evening, or in between.

###### CSD

All participants were requested to fill in a Consensus Sleep Diary (CSD; core form (7)) one week prior to the intake session, to obtain a general overview of their timing of sleep patterns.

###### SMH

To probe differences in state sleep quality, the St. Mary’s Hospital sleep questionnaire (SMH) was administered during all experimental sessions after sleep, as a systematic measure of the subjective quality of an individual’s previous night’s sleep (8).

##### Mood induction instructions

The instructions for the negative mood induction clips were, translated from the original Dutch instructions: *You will watch two clips from the movie “Sophie’s Choice”. The two clips will run after each other. It will last about 20 minutes in total. Try to sympathize with the main character. This is Sophie, the blonde-haired woman. Try to imagine how you would feel if you were in her situation. Let your emotions be influenced by the movie and try to maintain these*.

##### False memory task construction

To probe the activation of emotional schemas and measure emotional memory bias, we constructed new emotional semantically associated lists using the Dutch version of the Affective norms for English words (ANEW) list (9). The highest 85 negatively and 85 positively rated (on a 15-point Likert scale) and semantically independent words with a length ranging from five to thirteen letters (mean: 8.6 ± 2.2) were selected from the ANEW list. From these words, we selected 24 words each that would activate distinct emotional memory schemas. These words were chosen to function as the critical lure. To construct the lists of associations, 53 native Dutch participants participated in an initial pilot study and were asked to write down 10 associated words per presented word via an online questionnaire. The most frequently mentioned associates (excluding duplicates) were included in the final lists. Next, 24 independent native Dutch participants were asked to rate the lists of words. Specifically, they were asked to order the words according to the extent to which they thought the associations fitted the lure. These ratings determined the order of the final lists. After subsequent piloting (*N* = 50) of the false recall and recognition rates and timing presentation of each list, the 20 lists (10 per valence) of 10 words each that led to the highest numbers of false recall and recognition were included in the final version. The encoding session thus consisted of the presentation of a total of 200 words, corresponding to 20 lists of 10 words. From each list presented at encoding, the first, seventh and ninth word were included as old items, in correspondence to previous experiments, leading to a total of 60 old words. An additional 60 new words from the discarded lists and all 20 critical lures were included, resulting in a total of 140 words included in the recognition test.

##### Sleep recordings

Both mastoids (M1 and M2) were used as reference and the Cz location as online reference. A ground electrode was placed at location Fpz. One electrooculography (EOG) electrode was placed approximately 1 cm below the left outer canthus, and another was placed approximately 1 cm above the right outer canthus, both references to M1. To record nightly muscle activity, one electromyography (EMG) electrode was placed in the midline, 1 cm above the inferior edge of the mandible, and two EMG electrodes were placed below the inferior edge of the mandible (2 cm to the right and left of the midline). Two electrocardiography (ECG) electrodes were placed under the rib cage of the participant and one below the collarbone. Sleep data was filtered during recording by the SOMNOscreen™ as follows: EEG high- and low-pass filter with 0.3 Hz and 35 Hz, respectively (with ECG elimination); EOG high-pass filter 0.3 Hz, low-pass filter 35 Hz; EMG high-pass filter 10 Hz, low-pass filter 100 Hz, notch filter 50 Hz; ECG high-pass filter 0.3 Hz, low-pass filter 70 Hz, and a notch filter set to 50 Hz. Sleep scoring was performed according to AASM standards using the SpiSOP toolbox developed by Weber (2013). Epoch length was set to 30 seconds. During sleep scoring, movement artifacts were rejected, according to AASM guidelines.

##### MR neuronavigation procedure

Anatomical scans were either retrieved from the local database by subject ID or acquired prior to the intake session to obtain a personalized image for neuronavigation during the TMS sessions. Structural scans were acquired using a T1-weighted magnetization-prepared rapid acquisition gradient echo (MP-RAGE) sequence with the following parameters: TR = 2300 ms; TE = 3.03 ms; flip angle = 8°; FOV = 256 X 256 mm; voxel size = 1 mm isotropic.

##### TMS procedure

Our control rTMS (5Hz) protocol was based on the study by (53), and was employed to control for the peripheral and possibly unpleasant sensations induced by cTBS. Given the affective nature of our main task, we aimed to avoid a confound across our two experimental groups by differing levels of perceived discomfort which might occur when using a different stimulation site (e.g., the typical vertex control site is further removed from facial muscles than the mPFC stimulation site). Therefore the same target location (forehead) was used as a control condition, but with a different stimulation protocol. The study by (53) found no behavioral differences between a 5Hz-aPFC or cTBS-vertex control stimulations in a similar set-up as our study.
In both protocols, the stimulation intensity was anchored relative to 80 % of the measured active motor threshold from the tibialis anterior, given that the representational motor area for the tibialis anterior effector is located at a similar depth in the interhemisperic fissure to our target location of the medial prefrontal cortex. During the intake session, the active motor-evoked threshold (aMT) of the tibialis anterior as well as the first dorsal interosseous was determined, as measured by electromyographic (EMG) recordings in response to single TMS-pulses delivered to the appropriate motor hotspots in the primary motor cortex and following the method of limits (10, 11). The EMG-signal was measured at a 1 kHz sampling rate and bandpass filtered at 1-1000 Hz using an EKIDA DC amplifier (Ekida GmbH). The aMT was determined as the lowest stimulation intensity at which 5 out of 10 pulses evoked a visible motor evoked potential (MEP) in the EMG recordings. The intensity of the TMS stimulation was defined at 80% of the aMT of the tibialis anterior, unless this intensity fell above 120% of aMT of the first dorsal interosseous. In the latter case, to ensure that stimulation intensity was within established safety guidelines (12), the intensity was adjusted to fall below 120% of the active first dorsal interosseous motor threshold. Participants whose achieved stimulation intensity was lower than 70% of target intensity were replaced (*N* = 5). The TMS-intervention was delivered twice, once at the intake session to determine tolerability, and in the first experimental session as an experimental intervention. Precise coil position for the TMS stimulation was established using neuronavigation with TMS Navigator Software (Localite®) based on the anatomical MRI of each participant. The coil was positioned tangentially to the skull, and oriented with a 90° angle with reference to the sagittal midline. This orientation has been shown to induce an electrical field in the lateral-medial direction, perpendicular to the cortical layers in the medial wall of the prefrontal cortex (13). The exact Talairach coordinates of the coil position during stimulation was recorded for each session. Mean stimulation target site was calculated to be at MNI coordinate of x = 0.7, y = 70.58, z = 28.82, see Supplementary Figure 1. The TMS protocols were applied with a MagVenture MC-B70 Butterfly coil, connected to a MagPro-X100 stimulator (MagVenture, Farum, Denmark), using a biphasic pulse configuration.

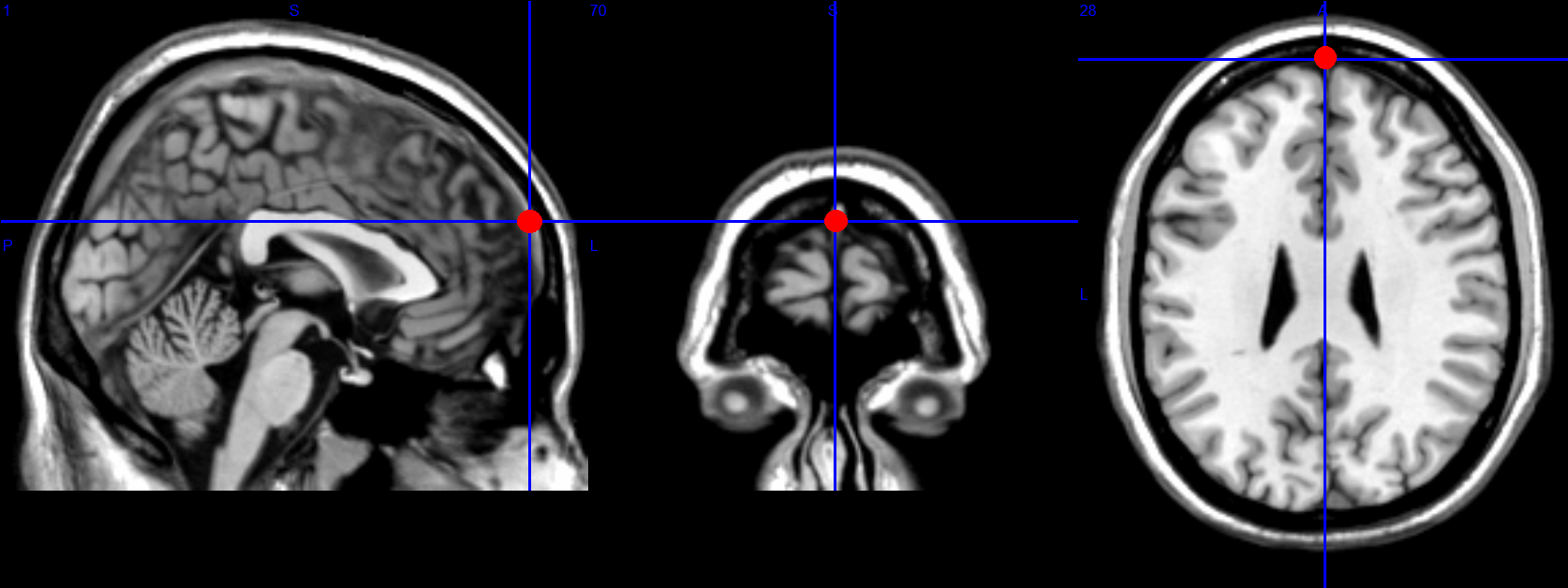

Supplementary Figure 1: Average stimulation site at the target region

#### Supplementary Results

##### Additional checks

The two experimental groups did not differ in their baseline measurements of depressive scores (BDI-I; *p* = 0.92), anxiety ratings (STAI-T; *p* = 0.69), neuroticism (NEO-PI-R; *p* = 0.32), circadian rhythm type (MEQ; *p* = 0.89) or general sleep quality (PSQI; *p* = 0.61). The groups also did not differ in their self-reported experience of discomfort from the two stimulation protocols, $t(37.65)=0.68$, $p=.503$. Furthermore, the groups did not differ in the “awareness” of the task procedure, $t(37.10)=0.22$, $p=.824$.

##### Mood induction results

There was a significant effect of time on both negative affect (NA; $F(2,112)=27.95$, $MSE=13.99$, $p<.001$) and positive affect (PA; $F(2,112)=3.79$, $MSE=33.45$, $p=.025$) scores, as well as their difference scores (NA-PA; $F(2,112)=20.58$, $MSE=43.87$, $p<.001$). See Supplementary Figure 2 depicting mood change in NA over time. A post hoc Tukey test showed that both groups show an increase in NA after negative mood induction, (*p* < .001). Scores returned back to baseline after having performed the encoding task, *p* < .001). There was no influence of NA differences between the timepoints before and after the movie, on the falsely remembering negative lures, $t(35)=-0.57$, $p=.571$, nor on the ratio of positive and negative lures, $t(35)=0.53$, $p=.600$. The interaction effect between group and time (before and after the movie) on the number of falsely remembered negative lures was also not significant, $t(35)=1.71$, $p=.096$.

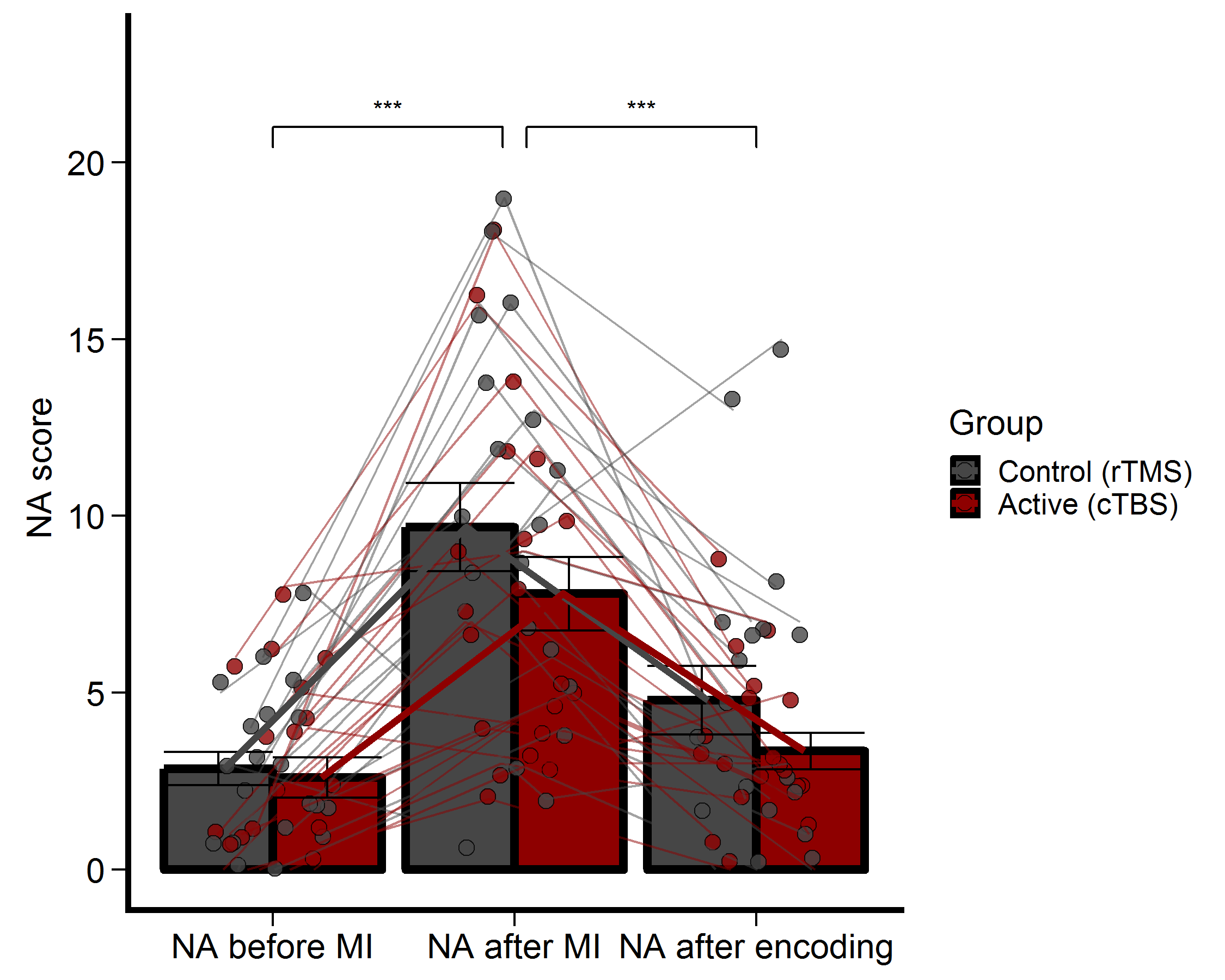

Supplementary Figure 2: Mood measurements. Negative moods were significantly increased after the negative mood induction. NA = Negative Affect, MI = Mood Induction.

##### Influence of depression and anxiety scores on memory bias

No influence of BDI scores $r^{2}$ = 0.00, *p* = 0.73 nor STAI scores $r^{2}$ = 0.03, *p* = 0.26 were found on the ratio of lures remembered (i.e. memory bias) across our participants.

##### Sleep data

The number of minutes in wake after sleep onset (WASO), a measure of sleep quality in the night between encoding and retrieval, displayed a negative relation to the proportion of correctly recognized old items using a Spearman correlation test, *r* = -0.33, *p* = 0.04 but not for the proportion of falsely recognized lures, *r* = -0.21, *p* = 0.20, see Supplementary Figure 3. The WASO did not influence the ratio of items recognized, $t(38)=-0.13$, $p=.901$. In contrast, subjective sleep quality did not influence memory performance of old items, $t(37)=0.06$, $p=.956$, nor of critical lures, $t(37)=0.05$, $p=.959$.

No significant results were found on the influence of WASO difference between the first and second night recording on memory performance of recognition of old items; *r* = -0.12, *p* = 0.52 nor for the proportion of falsely recognized lures, *r* = -0.13, *p* = 0.46. In addition, the difference score in WASO did not influence the ratio of items recognized, $t(31)=-0.16$, $p=.875$, neither did the differences in subjective sleep quality between the first and second night influence memory performance of old items, nor of critical lures, $t(37)=0.05$, $p=.959$.

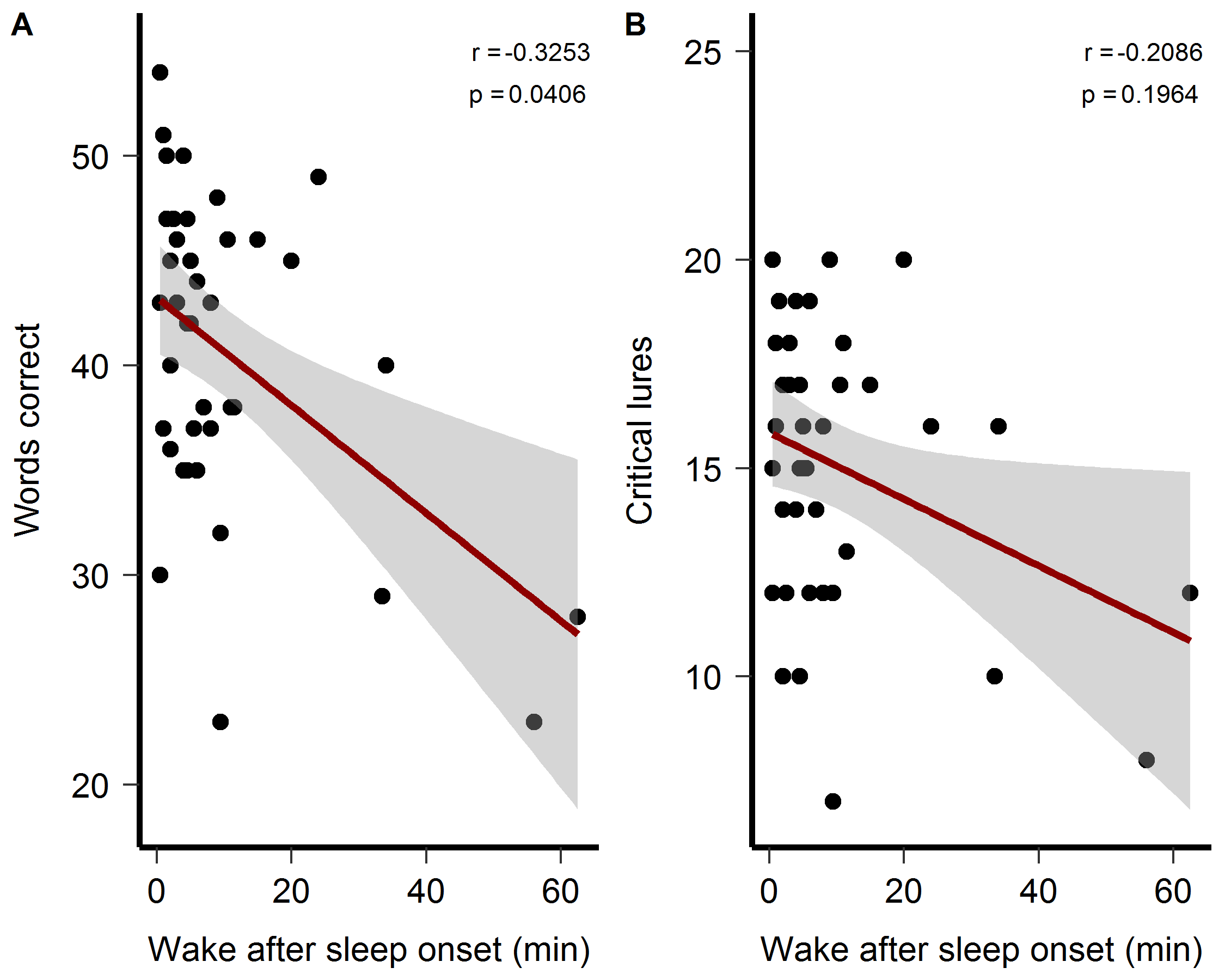

Supplementary Figure 3: Sleep quality and recognition memory. Sleep quality had an influence on veridical memory performance, but not false memory.

There was no influence of the number of time spent in stage 2 sleep during the second night on the recognition of old items, $t(38)=1.67$, $p=.103$ nor on the number of critical lures remembered,$t(38)=0.08$, $p=.933$. The influence of the difference in time spent in stage 2 sleep between the first and second night on the recognition of old items show similar results, $t(31)=0.81$, $p=.423$ as well as on the number of critical lures remembered,$t(31)=-0.47$, $p=.644$.

No influence of time spent in REM sleep on the ratio of valence of the critical lures was observed for the second night, $t(38)=-0.63$, $p=.532$, nor for the difference between the two recorded nights, $t(31)=-0.88$, $p=.387$. Since a few studies showed the influence of sleep spindles on emotional memory processing (14) as well as a relation with schema memory processing (15), we performed additional explorative analyses to look into the influence of different sleep spindle characteristics on false memory task performance. None of the fast spindle parameters during the second night were related to false memory recognition (spindle density: $r^{2}$ = 0.05, *p* = 0.18, count: $r^{2}$ = 0.03, *p* = 0.29, amplitude: $r^{2}$ = 0.01, *p* = 0.50 or duration, $r^{2}$ = 0.06, *p* = 0.14), nor ratio of lures (spindle density: $r^{2}$ = 0.00, *p* = 0.97, count: $r^{2}$ = 0.01, *p* = 0.61, amplitude: $r^{2}$ = 0.01, *p* = 0.64 or duration, $r^{2}$ = 0.01, *p* = 0.53). In turn, none of the fast spindle parameters were related to false memory recognition in the difference between the first and second night (spindle density: $r^{2}$ = 0.00, *p* = 1.00, count: $r^{2}$ = 0.00, *p* = 0.92, amplitude: $r^{2}$ = 0.04, *p* = 0.29 or duration, $r^{2}$ = 0.01, *p* = 0.58), nor ratio of lures (spindle density: $r^{2}$ = 0.01, *p* = 0.57, count: $r^{2}$ = 0.00, *p* = 0.77, amplitude: $r^{2}$ = 0.00, *p* = 0.81 or duration, $r^{2}$ = 0.01, *p* = 0.57).

As specific characteristics of REM sleep, especially REM density and REM latency, have been suggested as biomarkers for depression (16, 17), we explored if these parameters influence the ratio of memory for critical lures in our task. First, a partial Pearson correlation controlling for stimulation group, found no influence of REM latency in the night between encoding and retrieval on the ratio of critical lures remembered in all subjects combined, *r* = 0.11, *p* = 0.91, was there an influence of the difference between the first and second night, *r* = 0.94, *p* = 0.35. In addition, a similar analysis using REM density as predictor showed similar results, *r* = 0.27, *p* = 0.79.

##### Mood ratings during encoding

After each individual encoding list, the participants were asked to rate their change in mood, ranging from relatively sadder to relatively happier. The main purpose of the question was to ensure participants’ attention during encoding, but the collected data could additionally be analyzed to explore any association between mean mood rating and subsequent memory performance. Per list, 3 items were tested during recognition, thus a score of either 0, 1, 2 or 3 items correct could be assigned to each list per participant. No relation between mood changes and number of items correct was found, $t(798)=0.63$, $p=.526$.

##### Meta memory

After each list was presented, after the mood change question, a second question was asked, namely how many words each participant thought they would remember of that previous list (i.e. meta-memory). We tested the interaction between group and meta memory on performance of number of items correctly remembered using mixed effects modeling. To control for individual subject effects, a linear mixed-effects model was conducted with the lmer function (lme4 package), and subject identity was included as a random factor. P-values were determined by model comparison between a model that included the interaction term versus a model that excluded the interaction term. No significant interaction effect was found between stimulation group and meta memory score on the number of items correct, $\chi^{2}$ = 2.37, *p* = 0.31. A similar model was tested but this time including the false memory of the critical lure per list, thus a score of either 0 or 1, as a dependent variable. Here, a significant interaction effect was found between stimulation group and meta memory score on the number of falsely recognized critical lures, $\chi^{2}$ = 7.23, *p* = 0.03.
